# Integration of light quality signals regulates ABA abundance and stomatal movements during seedling establishment

**DOI:** 10.1101/2024.09.24.614678

**Authors:** Mathilda Gustavsson, Lionel Hill, Keara A. Franklin, Ashley J. Pridgeon

**Affiliations:** School of Biological Sciences, Life Sciences Building, University of Bristol, BS8 1TQ; John Innes Centre, Norwich Research Park, Norwich, NR4 7UH

## Abstract

Obtaining sufficient light for photosynthesis and avoiding desiccation are two key challenges faced by seedlings during early establishment. Perception of light quality via specialised photoreceptors signals the availability of sunlight for photosynthesis. Canopy shade is depleted in red (R) and enriched in far-red (FR) light, lowering R:FR ratio, while direct sunlight and sunflecks contain UV-B. The balance of these wavelengths can determine the developmental strategy adopted by seedlings to either avoid shade, via stem elongation, or promote the expansion of photosynthetic organs. How seedlings regulate stomatal movements in different light environments is poorly understood. Using FR and UV-B supplementation to mimic aspects of canopy shade and sunlight respectively, we monitored stomatal apertures in *Arabidopsis thaliana* cotyledons. We show that low R:FR inhibits stomatal opening via a mechanism involving PHYTOCHROME INTERACTING FACTOR 4 (PIF4) and increased abscisic acid (ABA). In contrast, UV-B perceived by the UV RESISTANCE LOCUS 8 (UVR8) photoreceptor antagonises this response to promote stomatal opening in a response requiring phototropin photoreceptors. The convergence of phytochrome and UVR8 signalling to control ABA abundance enables plants to coordinate stem elongation and water use during seedling establishment in dynamic light environments.

## Introduction

Plants have evolved a suite of photoreceptors to detect light quality and quantity, signalling the availability of sunlight to fuel photosynthesis. Seedlings emerging under canopy shade are exposed to increased amounts of FR light, reflected and transmitted from vegetative tissue. This lowers the ratio of red to far-red light (R:FR), inactivating phytochrome photoreceptors and driving stem elongation to overtop competitors ^1^. Direct sunlight and sunflecks contain UV-B light which antagonises shade avoidance responses ^2–4^, promoting a compact stature ^5^. Continuous monitoring of light quality changes therefore enables plants to optimise developmental strategy in fluctuating environments ^6^.

Mortality rates are high during early seedling establishment ^7^ with water scarcity presenting a significant stress ^8^. Stomata are microscopic pores found predominantly on the epidermis of leaves. They perform a crucial role in the prevention of water loss and the uptake of carbon through the regulation of gas exchange. This can involve short-term alterations in the size of the stomatal pore and longer-term alterations in stomatal development, changing the number of stomatal pores on a plants surface) ^9^.Stomata are sensitive to a range of exogenous and endogenous signals (including light, temperature, CO_2_ concentration, water-availability, pathogen and microbe associated signals, among others). The signalling processes involved in blue light-induced stomatal opening ^10^ and abscisic acid (ABA)-induced stomatal closure are well characterised ^9,11^. Although guard cells can respond autonomously to environmental signals such as blue light and ABA, responses can be modulated by mesophyll-produced signals such as sugar and malate to coordinate stomatal aperture with leaf carbon assimilation ^9,12^. Compared to the blue light response the stomatal opening response to red light is less well understood. As the red-light response saturates at similar light intensities to photosynthesis it is thought to be linked to photosynthetic carbon assimilation, however, the exact nature of this link is unclear ^13,14^. Additionally, studies have shown that the red light photoreceptor, phytochrome B (phyB) and downstream phytochrome signalling components contribute to the coordination of stomatal apertures ^15–17^.

In this study, we analysed how light quality signals controlling light foraging affect stomatal aperture in developing seedlings. Stomatal development is established during embryogenesis with stomata forming rapidly after germination ^18^. The stomata of young seedlings are responsive to light signals ^15^, but the role stomatal movements play during seedling establishment is poorly understood. Here, we analysed cotyledon stomatal apertures treated with white light (WL) supplemented with FR and/or low dose UV-B to simulate aspects of canopy shade and direct sunlight respectively. We show that low R:FR inhibits stomatal opening in cotyledons in a response involving the basic helix-loop-helix (bHLH) transcription factor, PIF4 and accumulation of ABA. In contrast, UV-B supplementation promotes the sustained opening of stomata in a process requiring the UVR8 and phototropin photoreceptors as well as the signalling component BLUE LIGHT SIGNALLING 1 (BLUS1). When low R:FR and UV-B signals are combined, UV-B signalling overrides the effects of low R:FR on ABA abundance and stomatal aperture. Our data suggest that the integration of light quality signals by multiple photoreceptors coordinates seedling light foraging strategy with water use through the alteration of hormone levels.

## Results

### Low dose UV-B enhances stomatal opening in a UVR8- and phototropin-dependent manner in seedling tissue

To assess the effect of low dose UV-B (1 µmol m^-2^s^-1^) on stomatal aperture, 7-day-old Arabidopsis seedlings were transferred to white light (WL) ± UV-B at dawn (light spectra are presented in **Fig. S1**). Cotyledon stomatal apertures were recorded over a 6 h period and are presented in **Fig. 1a**. In contrast to reports using older leaves and/or higher doses of UV-B or epidermal strips ^19–22^, seedlings supplemented with UV-B showed enhanced stomatal opening when compared to WL controls during the latter part of the time course. Exposing seedlings to monochromatic UV-B (**Fig. 1b**) showed that the UV-B-induced enhancement of stomatal opening requires a background of light within the 400-700nm wavelength range. Stomatal responses to UV-B were also analysed in mature plants. Here, apertures were measured from rosette leaves, and in leaf disc and epidermal peel tissue treated while floating on buffer. In contrast to cotyledons, no UV-B-enhanced stomatal opening was observed in rosette leaves (**Fig. S2a-c**). Some UV-B-mediated increase in stomatal aperture was, however, observed in leaf disc and epidermal tissue (**Fig. S2d-e**).

Blue light-induced stomatal opening involves the redundant actions of phototropin photoreceptors, phot1 and phot2 ^23^. Analyses of the photoreceptor mutants *uvr8-6* and *phot1/2* showed that both are required for UV-B-enhanced stomatal opening (**Fig. 1c and d**), suggesting a role for phototropin and UVR8 signalling. Additionally, mutants deficient in the downstream phototropin signalling component BLUS1 (*blus1-3*) ^24^ displayed a similar phenotype to *phot1/2* (**Fig. 1e**), with no additional opening in response to UV-B supplementation observed. These data provide further support for a role for phototropin signalling in UV-B-enhanced stomatal opening.

Following UV-B absorption, UVR8 dimers monomerise and interact with the E3 ubiquitin ligase CONSTITUTIVELY PHOTOMORPHOGENIC (COP1) ^25,26^. This stabilises the bZIP transcription factors ELONGATED HYPOCOTYL 5 (HY5) and HY5 HOMOLOGUE (HYH) which control the expression of a number of UV-B regulated genes ^27,28^. We further investigated the involvement of these components in the enhanced stomatal opening response observed under UV-B supplemented conditions. The *hy5/hyh* double mutant displayed wild type stomatal responses to UV-B light, suggesting that these transcription factors are not required for UV-B-mediated stomatal opening (**Fig. S3a**). This mutant is in the Wassilewskija (Ws) background, which behaved similarly to Col-0, confirming consistency of this response across different Arabidopis accessions. Consistent with previous reports ^29–31^, the *cop1* mutant showed consistently open stomata under all conditions, but did not respond to UV-B supplementation (**Fig. S3b**). Additionally, we observed that the *tt4* mutant (a flavonoid biosynthesis mutant hypersensitive to UV-B and more prone to UV-B induced damage ^9,32^) showed no UV-B-mediated enhancement of stomatal opening, suggesting damaging amounts of UV-B may prevent the enhanced opening we observe at 6 hours (**Fig. S3c**).

**Fig. 1.**
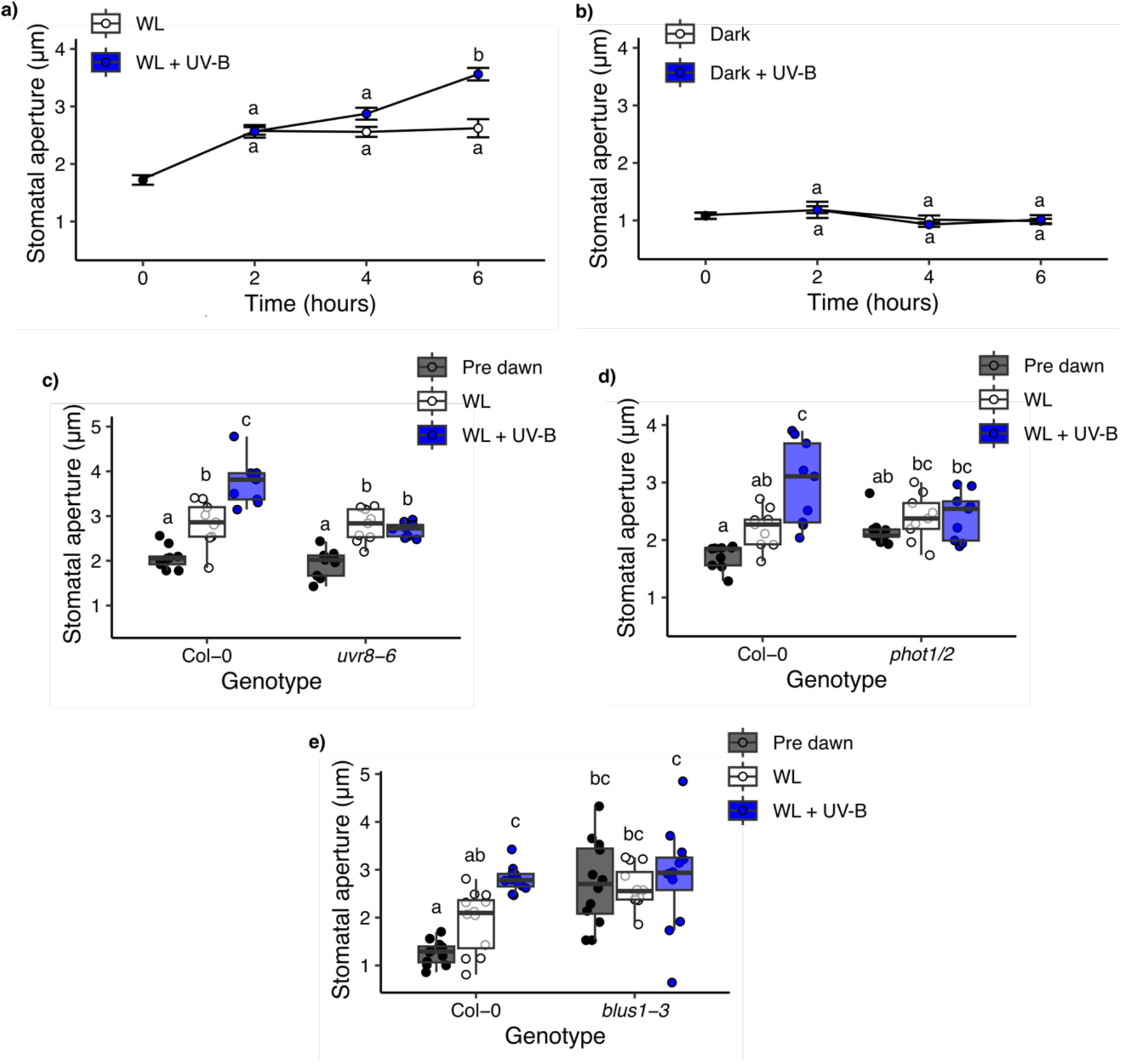
Low dose UV-B enhances stomatal opening in a UVR8- and phototropin-dependent manner. Stomatal apertures of 7-day-old Arabidopsis seedlings in response to different light treatments. Seedlings were treated with **a)** white light (WL) ± UV-B and **b)** darkness ± UV-B (Dark + UV-B) over a 6 h time course. Data are presented as the mean ± s.e. of each plant’s average stomatal aperture. The stomatal responses of **c)** *uvr8-6*, **d)** *phot1/2,* and **e)** *blus1-3* mutants following 6 h of WL ± UV-B treatment. Treatments were started at dawn and apertures measured prior to dawn (Pre dawn). Data are presented as boxplots showing the median and interquartile range of each group. The upper and lower whiskers represent data within 1.5 * IQR. Each individual plants mean stomatal aperture is represented as a point on the plot. For all genotype treatment combinations n = 9-12 seedlings over 3 independent experiments. Each mean seedling stomatal aperture was calculated from 10-12 stomatal measurements. Data were analysed using a 2-way ANOVA, followed by Tukey multiple comparison test.

### Low R:FR ratio treatment inhibits stomatal opening and is antagonised by UV-B

Reduced R:FR is a major component of vegetative shade which drives inactivation of phytochrome photoreceptors, auxin production and the elongation of stems for light foraging^1,6,33^. The effect of reduced R:FR (achieved by supplementing WL with FR) on seedling stomatal apertures was investigated (**Fig. 2a, Fig. S2**). Low R:FR decreased stomatal aperture in a response observed after 1-3 h of treatment. When WL was supplemented with both FR and UV-B light, stomatal apertures were no longer reduced, suggesting that the presence of UV-B overrides the effect of low R:FR. Decreased stomatal apertures were also observed in rosette leaves, although at later time points than in cotyledons (**Fig. S2a-c**).

The role of PIF transcription factors in stomatal responses to low R:FR was next explored, as PIFs have been shown to regulate both shade avoidance ^34–36^ and stomatal aperture ^15,16^. PIF4, PIF5 and PIF7 are the major regulators of plant architectural responses to low R:FR, so we therefore focused on these genes ^34,36–38^. *pif7* mutant seedlings displayed a wild-type response to reduced R:FR (**Fig. 2b**), whereas *pif4* and *pif4/5* mutants showed more variable stomatal apertures, with no significant differences between WL and low R:FR (**Fig. 2c and d**). Furthermore, the *pif4/7* double mutant displayed an insensitivity to low R:FR conditions with significantly more open stomata in low R:FR than Col-0 (**Fig 2d**). This suggests a dominant role for PIF4 in inhibiting stomatal opening under low R:FR conditions with a potential minor redundant role for PIF7.

PIF activity is regulated by the red and far-red light-absorbing phytochrome photoreceptors ^39^, with phyB performing a major role. In high R:FR, activated phyB binds to PIFs, leading to their ubiquitination and degradation via the 26S proteosome ^40–42^. The stomatal response of *phyB* mutants to low R:FR was therefore explored (**Fig. 2e**). *phyB* mutants displayed significantly smaller stomatal apertures than wild-type controls in white light which were not further decreased in low R:FR. Both These data suggest that phyB inactivation in low R:FR stabilises PIF4 which promotes stomatal closure.

To assess whether light treatments were indirectly affecting stomatal apertures through leaf temperature changes, thermal imaging was used to track cotyledon temperature over a 6 h time course (**Fig. S4a-b**). At 2 h post dawn there was no significant difference in cotyledon temperature between WL and WL + FR treatments. However, treatments involving the addition of UV-B showed cotyledons to be ∼0.5°C warmer than treatments without UV-B (**Fig. S4c**). Higher temperatures have been shown to elicit increased stomatal opening, however this effect has been reported at temperatures greater than 35 °C ^43–45^. To assess whether slightly elevated cotyledon temperature may account for the increased stomatal opening observed under UV-B supplemented conditions, seedlings were treated ± UV-B at 20 and 28 °C (**Fig. S4d**). No effect of raised temperature was observed in WL, and UV-B promoted enhanced opening at 20°C and 28°C. Additionally, the stomatal density of *uvr8-6* and *pif4* mutant cotyledons were analysed and found to show no significant differences from Col-0 (**Fig. S5**), suggesting the differences in UV-B and FR responses observed across these genotypes are not due to differences in stomatal density.

**Fig. 2.**
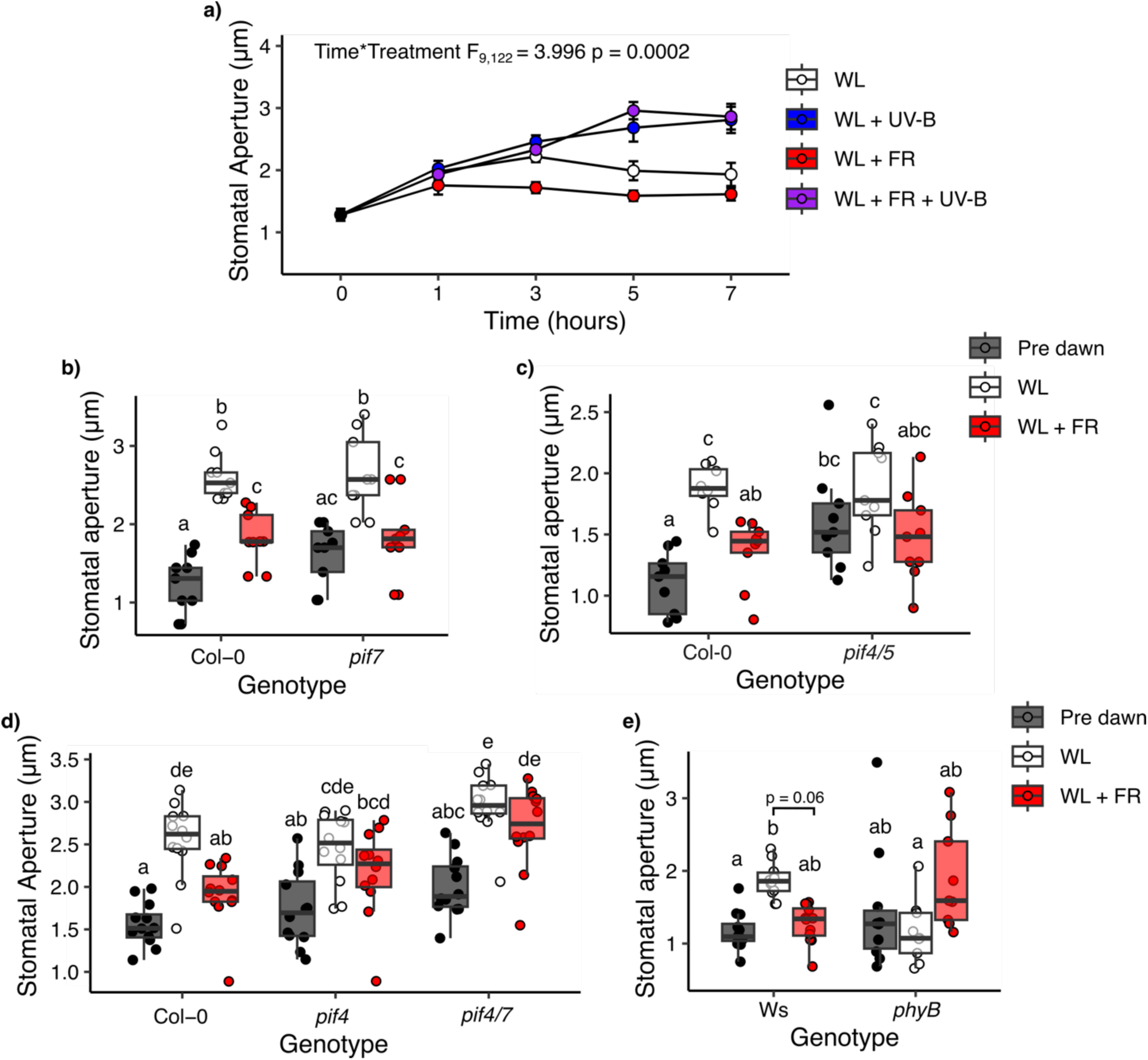
Low R:FR inhibits stomatal opening in a PIF-dependent manner and is antagonised by UV-B. **a)** Col-0 seedling stomatal apertures were monitored over a 7 h time course where plants were treated with white light (WL) ± far red (FR) and/or low dose UV-B. **b-c)** Col-0 and *pif* mutant seedlings were treated with WL ± FR for 2 h before measurement of stomatal apertures. These were also measured prior to dawn (Pre dawn). In **a)** data is presented as the mean ± s.e. of each plants average stomatal aperture. In **b-c)** data is presented as boxplots showing the median and interquartile range of each group. The upper and lower whiskers represent data within 1.5 * IQR. Each individual plant’s mean stomatal aperture is represented as a point on the plot. For all genotype treatment combinations n = 9-12 seedlings over 3 independent experiments. Each mean seedling stomatal aperture was calculated from 10-12 stomatal measurements. Data was analysed using a 2-way ANOVA, followed by Tukey multiple comparison test.

### R:FR and UV-B control ABA levels

Abscisic acid (ABA) performs a major role in regulating stomatal responses to drought stress. ABA is a potent trigger of stomatal closure and functions to keep stomata closed by inhibiting stomatal opening ^46,47^. Multiple studies have reported links between the biosynthesis and signalling of abscisic acid (ABA) and phytochrome signalling ^16,48–51^. Recently, PIF and ABA accumulation dynamics were observed to perform a role in the regulation of stomatal apertures over day/night cycles ^15^. ABA signalling has also been linked to UV-B responses as, in *Zea mays*, high dose UV-B treatment stimulates ABA production ^52^.

To investigate the role of ABA signalling and/or biosynthesis in stomatal responses to UV-B and reduced R:FR, we assayed the ABA content of Col-0, *uvr8-6,* and Ws seedlings treated with UV-B, FR, or a combination of the two light treatments after 6 h. Analysis of Col-0 and *uvr8-6* data in **Fig. 3a** via 2-way ANOVA shows a significant effect of both genotype (F_1,32_ = 8.15, p = 0.008) and light treatment (F_3,32_ = 11.45, p < 0.001). In **Fig. 3b** analysis of Ws via 1-way ANOVA shows a similar significant effect of light treatment (F_3,8_ = 14.78, p = 0.001). In both Col-0 and Ws accessions, increased ABA content was repeatedly observed in low R:FR conditions, whereas decreased ABA was observed in the presence of UV-B. When low R:FR and UV-B treatments were combined, an intermediate ABA content was observed, resembling levels observed in WL. Together, these data suggest that UV-B perceived by UVR8 can counteract low R:FR-induced increases in seedling ABA (**Fig. 3a**).

To investigate the potential role of ABA signalling further, stomatal responses of mutants defective in ABA biosynthesis (*nced3/5* – defective in *de novo* ABA biosynthesis ^53^, and *bg1 –*deficient in an enzyme that rapidly generates active ABA from a pool of inactive ABA glucosyl-ester (ABA-GE) ^54^) and ABA signalling (*q1124 –* a quadruple ABA receptor mutant ^55^ and *ost1-3 –* a mutant in a key downstream kinase ^56^) were assayed in response to UV-B and low R:FR treatments. Here, the ABA biosynthesis mutants *nced3/5* and *bg1*, and the quadruple receptor mutant *q1124* displayed wild-type responses to UV-B supplementation, whereas the *ost1-3* mutant showed no significant increase in stomatal opening (**Fig. 3c-e**). The *nced3/5* ABA biosynthesis and *q1124* ABA receptor mutants displayed increased apertures in WL, supporting previous observations ^57,58^. In all ABA biosynthesis and receptor mutants, no significant differences were observed between WL and low R:FR conditions, supporting the involvement of ABA in the low R:FR-mediated inhibition of stomatal opening (Fig 3f). Interestingly, this phenotype was most pronounced in the *bg1* ABA biosynthesis mutant, suggesting that deconjugation of ABA from ABA-GE is central to elevating ABA levels in low R:FR conditions (**Fig. 3f**). Similarly to *bg1*, the *ost1-3* signalling mutant showed no stomatal aperture response to low R:FR conditions, confirming the importance of OST1 in stomatal aperture regulation ^56^ (**Fig. 3g**). Together, these data suggest that ABA biosynthesis and signalling are not essential for UV-B-induced stomatal opening but are required for low R:FR-induced inhibition of stomatal opening.

**Fig. 3.**
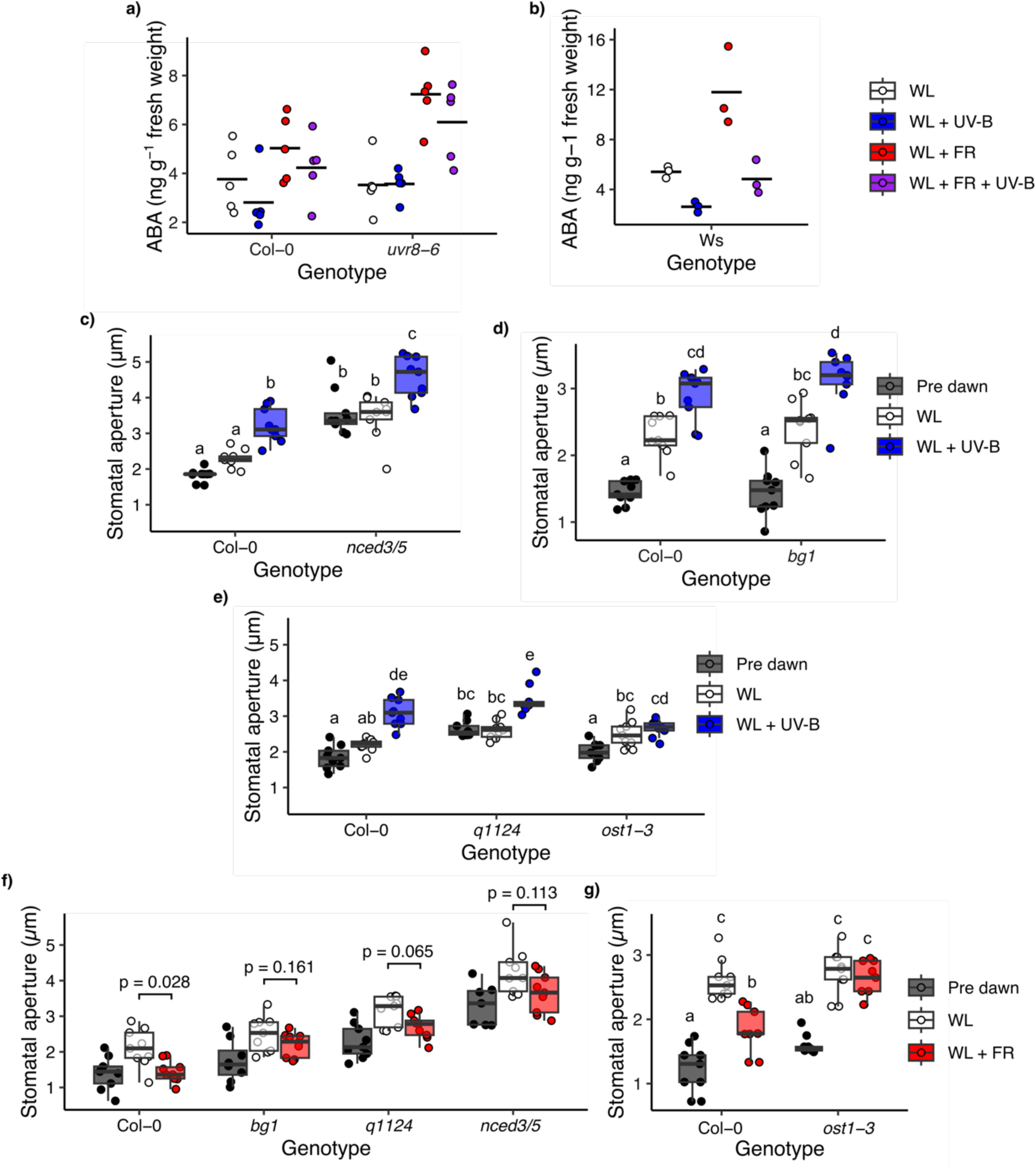
Low R:FR promotes ABA accumulation via a mechanism inhibited by UV-B. ABA levels in the aerial tissue of 7-day-old **a)** Col-0 and *uvr8-6,* and **b)** Ws seedlings treated for 6 h with white light (WL) ± far red (FR) and/or UV-B light and quantified using LC-MS. Stomatal apertures of 7-day-old **c)** ABA biosynthesis mutants *nced3/5* and **d)** *bg1,* and **e)** ABA signalling mutants *q1124* and *ost1-3,* were measured predawn and following a 6 h WL ± UV-B treatment. The stomatal aperture response of ABA biosynthesis and signalling mutants **f)** *bg1*, *q1124*, *nced3/5*, and **g)** *ost1-3* were measured predawn and in response to 2 h WL ± FR treatment. For **a-b)** all data is presented as points and the mean as a horizontal bar. For all other plots, data are presented as boxplots showing the median and interquartile range (IQR) of each group. The upper and lower whiskers represent data within 1.5 * IQR. Each individual plant’s mean stomatal aperture is represented as a point on the plot. For all genotype and treatment combinations, **a)** n = 5 seedlings over 5 independent experiments and **b)** n = 3 seedlings over 3 independent experiments. For **c-g)** all genotype and treatment combinations, n = 9 seedlings over 3 independent experiments. Each mean seedling stomatal aperture was calculated from 10 stomatal measurements. All data were analysed using a 2-way ANOVA, followed by Tukey multiple comparison tests, except for **b)** where a 1-way ANOVA was used and **f)** where T tests were used to make specific comparisons and adjusted for multiple comparisons using a Holm correction. In **g)** *ost1-3* was grown in parallel with *pif7-2* (Fig. 2c) each plot uses the same Col-0 control data.

## Discussion

This study shows that the stomatal movements of seedlings are modulated by integrated light quality signals. White light supplemented with low dose UV-B promotes the additional opening of stomata (**Fig. 1, 2a**), whereas low R:FR acts oppositely to limit stomatal aperture (**Fig. 2**). Our data suggest that low R:FR inhibits stomatal opening through promoting ABA accumulation, and that UV-B light can antagonise this, in part, through suppression of ABA levels (**Fig. 3**).

### Low dose UV-B promotes stomatal opening in seedling cotyledons

In contrast to our observations, most studies focusing on stomatal responses to UV-B have shown UV-B irradiation to induce stomatal closure. A number of components underlying this response have been identified, including the UVR8 photoreceptor, COP1, HY5 and HYH ^19,20^, a G alpha protein ^21^, MAP KINASES (MAPKs ^22^, ethylene^20,59^, and Hydrogen Peroxide (H_2_O_2_) and Nitric Oxide (NO) ^21,60^. The latter are stress signalling components, most likely induced by the high doses applied.

Here, in contrast, we found that lower dose UV-B (1 µmol m^-2^ s^-1^) applied in a background of white light increases stomatal apertures of the cotyledon abaxial epidermis (**Fig. 1, 2a**). This occurs in both Col-0 and Ws Arabidopsis accessions (**Fig. 1, Fig. S3**), to a lesser extent in detached mature plant tissue, but not in non-detached mature plant tissue (**Fig. S2**). It is unlikely that the light treatments used in this study induced stress signalling, as Fv/Fm measurements of chlorophyll fluorescence (commonly used as an indicator of photosystem health ^61^) varied little in response to low R:FR or UV-B. Unsurprisingly, a mild reduction in Fv/Fm was observed in UV-B-treated *uvr8-6* mutants which are unable to initiate photoprotective responses (**Fig. S6**). Some studies have reported stomatal opening in response to UV-B supplementation in *Vicia faba* ^62^, cucumber ^63^ and certain Ericaceae species ^64^ and it has been suggested that the effect of UV-B light on stomata is dependent on the plant metabolic state ^62,65^.

We show that the opening of seedling abaxial stomata following UV-B treatment requires the presence of light within the photosynthetically active range (400-700 nm), functional phototropin and UVR8 photoreceptors (**Fig. 1**), together with the downstream phototropin signalling component BLUS1 (**Fig. 1e**). The requirement for phototropin signalling differs from observations showing that low doses of UV-B alone could stimulate stomatal opening in both Arabidopsis and *Vicia faba* ^66^. However, there are numerous differences between the plant growth conditions, plant ages, and experimental procedures that may explain the contrasting observations. In this study we have focused on the stomatal apertures of intact seedling cotyledon tissue and tracked apertures over a longer period through the course of the day.

Our data suggest that UV-B-mediated promotion of stomatal opening does not require the UV-B signalling components HY5 or HYH (**Fig. S3a**). This is in contrast to the UVR8-mediated promotion of stomatal closure ^19,20^. Upon assaying mutants deficient in the UV-B signalling component, COP1, we observed a constitutively open response before dawn, and no difference between WL and WL + UV-B conditions (**Fig. S3b**). These mutants have previously been shown to present a constitutively open stomata phenotype under dark conditions ^17,29–31^, confounding interpretation of the role of COP1 in UV-B-promotion of stomatal opening.

### Low R:FR inhibits cotyledon stomatal opening in a PIF4-dependent manner

We further show that seedlings exposed to low R:FR ratio display reduced stomatal apertures following 1-3 hours of treatment in a response requiring phyB and PIF4 (**Fig. 2, 3, S2**). Previous studies have shown low R:FR to inhibit stomatal opening in *Commelina communis* and orchids ^67,68^, with no effect observed in Arabidopsis ^69^. The differences in our observations may be due to using intact cotyledon tissue, as opposed to epidermal peels from mature leaves. More recently, monochromatic FR treatment has been shown to reduce stomatal apertures in rice in a mechanism requiring the PIF-homologue, OsPIL15 ^16^. PIFs have also been shown to play a role in regulating the daily rhythmic opening and closing of stomata ^15^. Rovira et al. (2024) show PIFs to act in opposition to ABA, promoting the accumulation of K+ import channel POTASSIUM CHANNEL IN ARABIDOPSIS THALIANA (KAT1) and ultimately stomatal opening at dawn in 3-day-old Arabidopsis seedlings. This positive role contrasts with the negative role identified in this study. This may reflect differences in plant age or growth conditions. Here, seedlings were soil grown, whereas Rovira et al used ½ MS plates, where humidity would be considerably higher. Growth under high humidity conditions is known to affect stomatal responses to several signals, including ABA, which may alter PIF signalling ^70^.

Several studies have identified connections between ABA signalling, phytochrome signalling, and red/far-red light responses. PIFs have been shown to bind to the promoter regions of ABA biosynthesis and signalling genes ^48^, as well as interact with the ABA receptor proteins PYL8 and PYL9 ^49^. Mutants deficient in phyB show increased ABA levels but reduced ABA sensitivity under well-watered conditions ^50^. ABA content has been shown to decrease in plants treated with red light ^71^, and conversely, increase (in a number of tissues including leaves and auxiliary buds) when plants are treated with low R:FR ^51,72–76^. Additionally, genes tagged with the GO term response to ABA are upregulated in low R:FR-treated Arabidopsis seedlings ^77^ and FR-treated leaf tips ^78^.

Here we observed multiple ABA signalling and metabolism mutants to show reduced responses to low R:FR, with the strongest phenotypes observed in *bg1* and *ost1-*3 (**Fig. 3**). These data support a mechanism whereby, in low R:FR, BG1 functions to generate active ABA from a pool of inactive ABA-GE, which may then, through ABA signalling, lead to the activation of the OST1 kinase, and ultimately the inhibition of stomatal opening.

### UV-B antagonises ABA accumulation and stomatal responses to low R:FR

When seedlings were exposed to both FR and UV-B supplementation simultaneously, UV-B antagonised low R:FR-mediated inhibition of stomatal opening (**Fig. 2a**). A parallel response is observed in hypocotyl elongation, where low R:FR-induced elongation is inhibited in the presence of UV-B. This results, in part, from UVR8-mediated sequestration of COP1, destabilising PIF proteins, independently from HY5/HYH-mediated signalling pathways ^2,3^. The addition of UV-B to low R:FR also affected seedling ABA content, reversing the low R:FR-stimulated increase back to WL levels (**Fig. 3a, b**). This, coupled with the lack of opening inhibition in ABA biosynthesis and signalling mutants (**Fig. 3f-g**) suggests an increase in ABA content is required for the low R:FR-mediated inhibition of opening. A hypothetical model outlining the mechanism behind this response is proposed in **Fig. 4**. In vegetational shade, low R:FR inactivates phyB, stabilising and promoting PIF4 activity. ABA accumulates, via *de novo* synthesis and de-conjugation of ABA-GE. When plants reach a gap in the canopy, UV-B perceived by UVR8 targets PIF4 for inactivation and degradation ^2,3^. ABA accumulation does not occur and repression of stomatal opening is removed. However, in non-shade (high R:FR) conditions, UV-B supplementation decreases ABA content in a UVR8 dependent manner (**Fig. 3b**).

### Conclusion: Seedlings integrate light quality signals to coordinate growth and stomatal aperture in dynamic light environments, via regulation of hormone abundance

Following germination, seedlings must balance growth and water use to transition to photoautotrophic development without depleting available resources. Light quality provides key information concerning the prevailing levels of vegetational shade and sunlight availability ^6^. Plants perceive light quality using multiple photoreceptors and integrate this information to regulate levels of multiple hormones, altering their physiology and development to optimise survival ^79^. Low R:FR and UV-B control auxin and gibberellin abundance and signalling to adjust plant architecture for maximum light capture. Here we show that low R:FR and UV-B also regulate ABA content in seedling aerial tissue to control stomatal movements. When a seedling emerges under a vegetative canopy, low R:FR-mediated inhibition of stomatal opening may allow seedlings to restrict water loss without greatly impacting photosynthetic rates. While testing this possibility is beyond the scope of the current work it is clear that low R:FR-mediated inhibition of opening would be overridden in the presence of low amounts of UV-B light, allow seedling stomata to rapidly respond to sun flecks (gaps in the canopy known to contain quantities of UV-B light) ^5^. When seedlings emerge from a canopy, they are exposed to increased amounts of photosynthetic light. R:FR increases, and the inhibition of stomatal opening is attenuated. Prolonged exposure to ambient UV-B would then further open stomata, ensuring maximum photosynthetic productivity in sunlight.

**Fig. 4.**
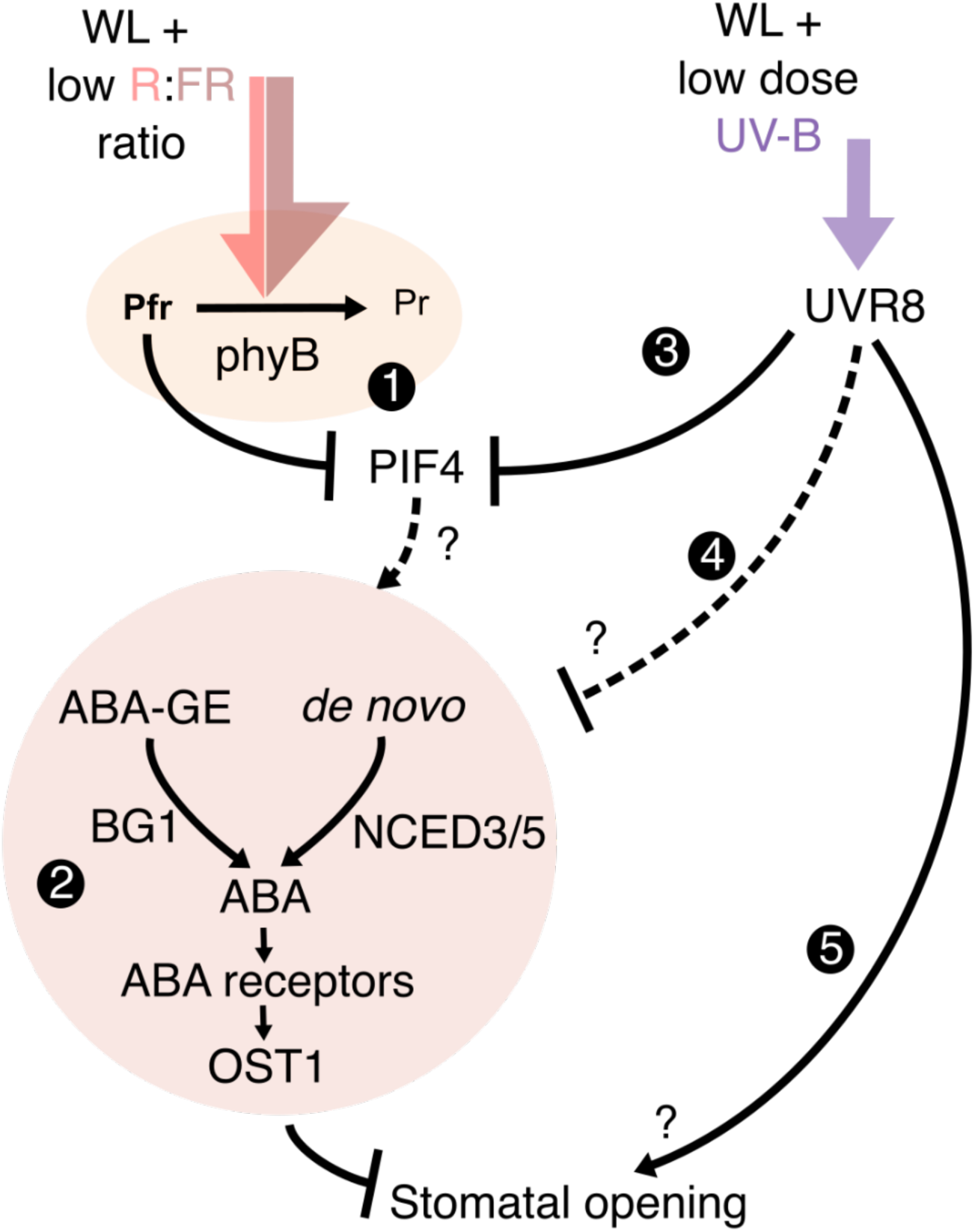
Proposed model for seedling stomatal aperture control by low R:FR and UV-B. A proposed mechanism for the action of low R:FR and UV-B on stomatal opening. **1)** Low R:FR promotes the conversion of the phytochrome B photoreceptor to its inactive Pr state, which, in turn, stabilises the phytochrome interacting factors (PIFs). In this study we observe PIF4 is required for seedling stomatal responses to FR supplementation. **2)** ABA biosynthesis and signalling components are required for inhibition of stomatal opening, likely downstream of PIF activity. The beta glucosidase enzyme (BG1) is involved in cleaving the glucose ester off an inactive pool of ABA-GE to rapidly generate active ABA and the NCED3 and NCED5 enzymes catalyse a key step in *de novo* ABA biosynthesis. Here we suggest that low R:FR-induced increases in seedling ABA levels drive inhibition of stomatal opening. UV-B supplementation abolishes the inhibition of stomatal opening in low R:FR. **3)** UV-B, perceived by UVR8 targets PIFs for inactivation and/or degradation. Inhibition of PIF function likely prevents the accumulation of ABA, relieving inhibition of stomatal opening. **4)** UVR8 may additionally function to directly inhibit ABA biosynthesis and/or signalling. **5)** UVR8 also promotes further stomatal opening through additional mechanisms unrelated to ABA signalling.

## Methods

### Plant material and growth conditions

*Arabidopsis thaliana* seedlings were grown in a 3:1 Levingtons F2 compost: silver sand (Melcourt) mixture. Seeds were washed in 70% ethanol before sowing and stratified for 2-3 days in the dark at 4 °C. Seedlings were grown under long day conditions (16 h day, 8 h night) grown in 75 µmol m^-2^ s^-1^ white light (see **Fig. S1** for light spectra) at a constant humidity and temperature of 70% and 20 °C respectively in growth cabinets (Microclima 1600E, Snijder Scientific, The Netherlands). All genotypes are in a Col-0 background apart from the *hy5hyh,* and *phyB* mutants which are in a Ws background. A list of genotypes ^26,53–56,80–87^ and their descriptions can be found in **Table S1**.

### Light treatments

Light measurements were performed using an Ocean Optics FLAME-S-UV–VIS spectrometer with a cosine corrector. For UV-B treatments a Philips TL100W/01 narrow band tube light wrapped in strips of heatproof tape was used to provide treatments of 1 µmol m^-2^ s^-1^ UV-B light (between 280 and 315 nm). Other than in **Fig. 1b**, UV-B was applied in a background of 75 µmol m^-2^ s^-1^ white light. For FR supplementation, LEDs emitting at a max peak of 735 nm were used in conjunction with 75 µmol m^-2^ s^-1^ white light. R:FR ratios were determined by dividing R light photon irradiance (between 660 – 670 nm) with FR light photon irradiance (between 725 - 735 nm). Low R:FR conditions were set to between 0.06-0.08 whereas high R:FR conditions were between 4-7.

### Stomatal aperture measurements

Arabidopsis seedlings were grown for 7 days under long day (16 h light, 8 h dark) conditions and transferred at dawn on the 8^th^ day to the appropriate light treatment unless otherwise stated. Length of light treatment is indicated in text and in each figure. The aerial portion of the seedling was rapidly placed on a microscope slide and images were taken of the abaxial surface of the cotyledons using a x40 lens on a Zeiss Axiovert 200M inverted microscope fitted with a Hamamatsu ORCA-ER digital camera. Images were randomised and stomatal apertures were then measured using FIJI (ImageJ) ^88^. For each experiment, stomatal apertures were measured predawn in addition to control and light treatments. Each experiment was repeated independently at least 3 times, with 3-4 plants measured per genotype per treatment. A total of 10-12 stomatal apertures were measured per plant, and the average of these stomatal aperture values was considered n = 1. Overall, each experiment has n = 9-12 for each genotype and treatment combination unless otherwise stated.

For **Fig. S2d,e,** leaf discs and epidermal peels were harvested from 4-5 week old plants grown under the same conditions as previously described for seedlings. Leaf discs were harvested using a 4 mm biopsy punch. After tissue harvesting, leaf discs and epidermal peels were transferred to 50 mm petri dishes containing 10 ml 10/50 buffer (10 mM MES, 50 mM KCl, pH adjusted to 6.15 using KOH) prewarmed to 20 °C. Dishes were incubated in darkness for 2 hours before measuring (pre-dawn) or transfer to 75 µmol m^-2^s^-1^ WL or WL supplemented with 1 µmol m^-2^s^-1^ UV-B. Leaf discs were measured 2, 4, and 6 h after treatment. Epidermal peels were measured 6 h after treatment. Each experiment was repeated independently at least 3 times, with 3 plants measured per genotype per treatment. A total of 10 stomatal apertures were measured per plant, and the average of these stomatal aperture values was considered n = 1. Overall, each experiment has n = 9 for each genotype and treatment combination.

### Stomatal density measurements

Arabidopsis seedlings were grown for 7 days under long day (16 h light, 8 h dark) conditions. On the 8^th^ day cotyledons were harvested and cleared ^89^. Cleared tissue was imaged using a x20 DIC lens on a Leica DMIRE2 microscope with a Leica DFC350 FX camera. A stack of images containing both the abaxial and adaxial epidermises was taken. Stomata were counted using the cell counter plugin of FIJI (ImageJ) within a 0.208 mm^2^ region.

### ABA quantification

Arabidopsis seedlings were grown and treated similarly to stomatal aperture experiments except after light treatment, aerial seedling tissues were rapidly harvested, the fresh weight was determined, and flash frozen in liquid nitrogen. Seedling tissue was ground into a fine powder and resuspended in 1.9 ml extraction buffer (1% v/v acetic acid in 100% isopropanol) spiked with 10 µl d6-ABA (2.5 µg ml^-1^ in 100% MeOH). Samples were left shaking overnight at 4 °C. These were centrifuged at 13.4 krcf for 5 mins at 4°C. Supernatant was evaporated 950 µl at a time into a new tube using an Eppendorf Concentrator Plus set to V-AL mode at 45°C for 1 hour. The original sample tubes were resuspended in 950 µl of extraction buffer (without d6-ABA) and left to shake for a further hour at 4 °C. The sample was centrifuged again at 13.4 krcf at 4 °C and the supernatant transferred to the tube containing the evaporated sample. This was evaporated a final time using the same settings as before. Dried samples were stored at −70 °C until mass spectrometry analysis.

Dried samples were resuspended in 100 µl 100% MeOH and run on an Acquity UPLC equipped with a XevoTQS tandem mass spec (Waters). Separation was on a 50×2.1 mm 2.6 μ Kinetex EVO C18 column (Phenomenex) using the following gradient of acetonitrile versus 0.1% formic acid in water, run at 0.7 ml min^-1^ and 30 °C (0 mins – 5%, 3 mins – 95 %, 3.5 mins – 95%, 3.6 mins – 5%, 5.1 mins – 5%). The samples were maintained at 10°C, and the instrument injected 5 μl. The hormones were detected by negative mode electrospray, with the following mass transitions: ABA, 263>153; d6-ABA, 269>159 (collision energy 10V). Spray chamber conditions were 900 l hr^-1^ drying gas at 500 °C, 150 l hr^-1^ cone gas, 7.0 bar nebulizer pressure, and a spray voltage of 1.5 kV.

### Thermal imaging

Thermal images were taken using a FLIR A665sc thermal imaging camera. ResarchIR software (v4.40.9.30) was used to generate TIF images which were then analysed using FIJI (ImageJ). Cotyledon temperatures were analysed within a 24px box and the mean of the 24 pixels was used as each cotyledons temperature value.

### Chlorophyll fluorescence

Chlorophyll fluorescence parameters of 7 day old seedlings were measured using a Imaging PAM M series system with ImagingWin software (v2.56p, Walz). Seedlings were treated with WL ± UV-B and ± FR for 6 hours. Following this, seedlings were dark adapted for 30 mins before applying a saturating pulse of light. Maximal photosystem II efficiency (F_V_/F_M_) was calculated using the formula F_V_/F_M_ = (F_M_ – F_0_) / F_M_. Where F_0_ and F_M_ represent fluorescence measurements before and after the saturating light pulse respectively. Images were exported from the ImagingWin software as .tif files and analysed using FIJI (ImageJ). Manual thresholding was used to generate masks of the seedling aerial tissue before calculating the mean F_V_/F_M_ for that region of interest.

### Data presentation and analysis

All data was statistically analysed in R (version 4.3.1) ^90^ and plotted using the ggplot2 package ^91^.

### Statistical Analysis

Data was analysed using R (version 4.3.1). Most datasets were analysed using 1 or 2-way ANOVAs with post hoc Tukey multiple comparison tests. Some datasets were analysed using Holm corrected T-tests.

## Acknowledgements

This work was supported by the Biotechnology and Biological Sciences Research Council-funded South West Biosciences Doctoral Training Partnership [training grant reference: BB/M009122/1], BBSRC grant BB/R002045/1, The Leverhulme Trust Early Career Fellowship [award number ECF-2022-310], and the Bristol Centre for Agricultural Innovation (BCAI). The authors thank Prof Antony Dodd for providing bench space at the John Innes Centre and Prof Alistair Hetherington for providing feedback on the manuscript.

## Author Contributions

M.G., K.A.F. and A.J.P. designed experiments. M.G., L.H., and A.J.P. performed experiments and analysed data. M.G., K.A.F., and A.J.P. wrote the manuscript.

## Supplementary Information

**Fig. S1.**
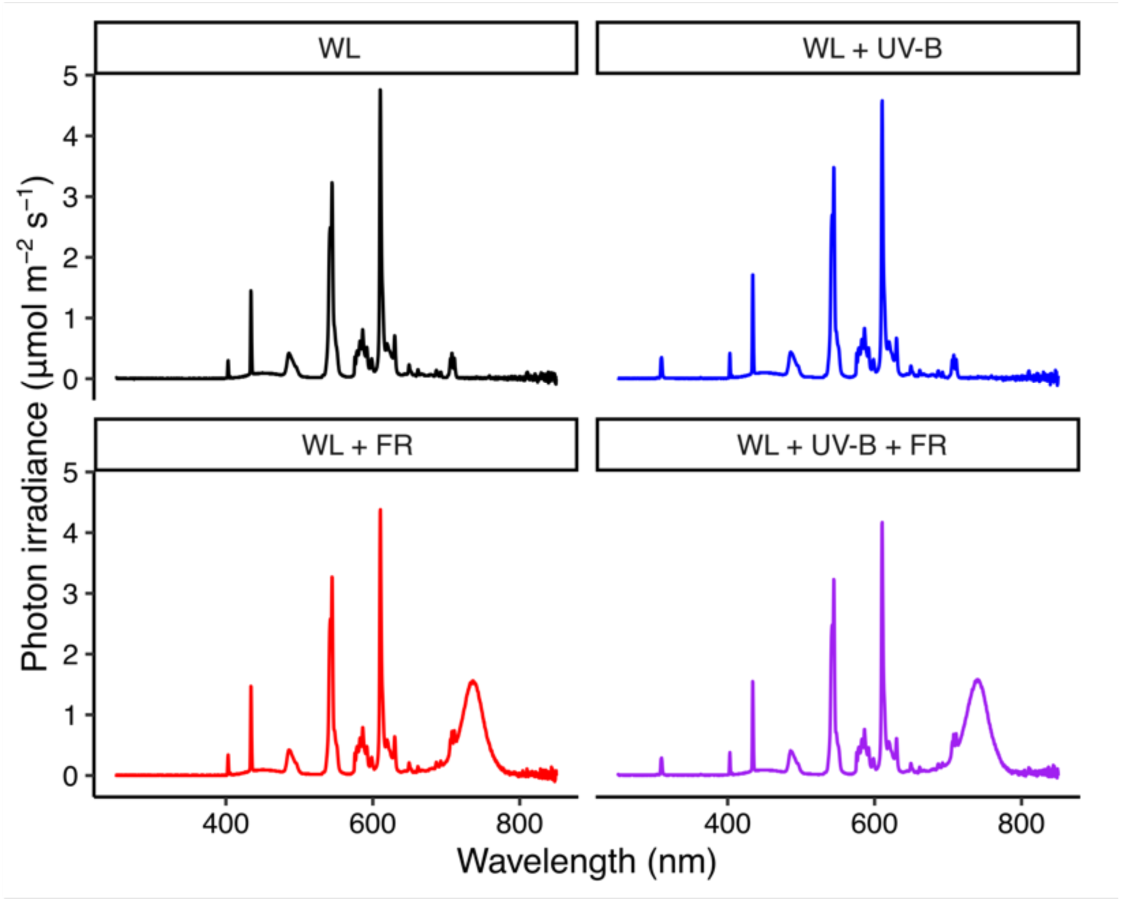
Cabinet spectrums. Light spectra of the different treatments described in this study. Every treatment includes ∼75 µmolm^-2^s^-1^ white light (light between 400-700 nm). UV-B treatments involved the addition of 1 µmolm^-2^s^-1^ of UV-B light provided by a Philips TL100W/01 narrow band tube light. FR treatments involved the addition of FR LEDs with a maximum emission peak at 735 nm.

**Fig. S2.**
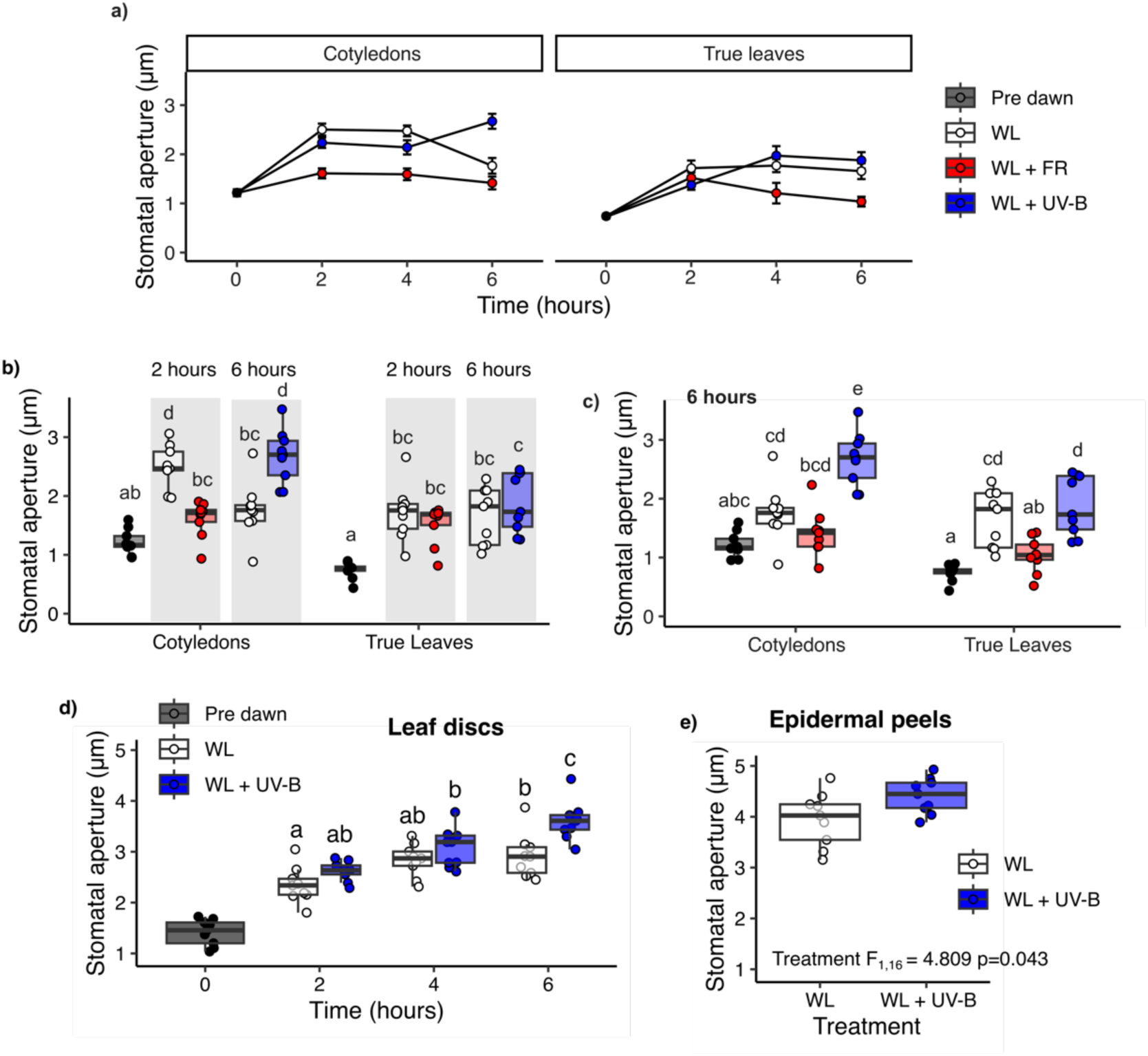
Cotyledons and rosette leaves differ in stomatal responses to light quality. **a)** Stomatal apertures from 7-day-old seedling cotyledons and 21-day-old true leaves were measured in WL, WL ± FR, and WL ± UV-B conditions over a 6 hour time course. Selected timepoints from **a)** are shown in **b)** 2h WL ± FR and 6h WL ± UV-B and **c)** all light treatments at 6 hours. Stomatal apertures from **d)** leaf discs and **e)** epidermal peels of 4-5 week-old Col-0 plants. Apertures were measured **d)** pre dawn and over a 6 h time course (2, 4, and 6 h) of WL ± UV-B treatment. In **e)** apertures were measured following 6 hours of WL ± UV-B treatment. For **a)** data is represented as mean ± s.e.m. For **b-e)**, data are presented as boxplots showing the median and interquartile range (IQR) of each group. The upper and lower whiskers represent data within 1.5 * IQR. Each individual plant’s mean stomatal aperture calculated from 10-12 stomatal measurements is represented as a point on the plot. For all genotype treatment combinations, n = 9 plants over 3 independent experiments. Data in **b-d)** were analysed using 2-way ANOVA followed by Tukey Multiple Comparisons tests. Data in **e)** was analysed using a Students T test. Letters indicate significance at p < 0.05.

**Fig. S3.**
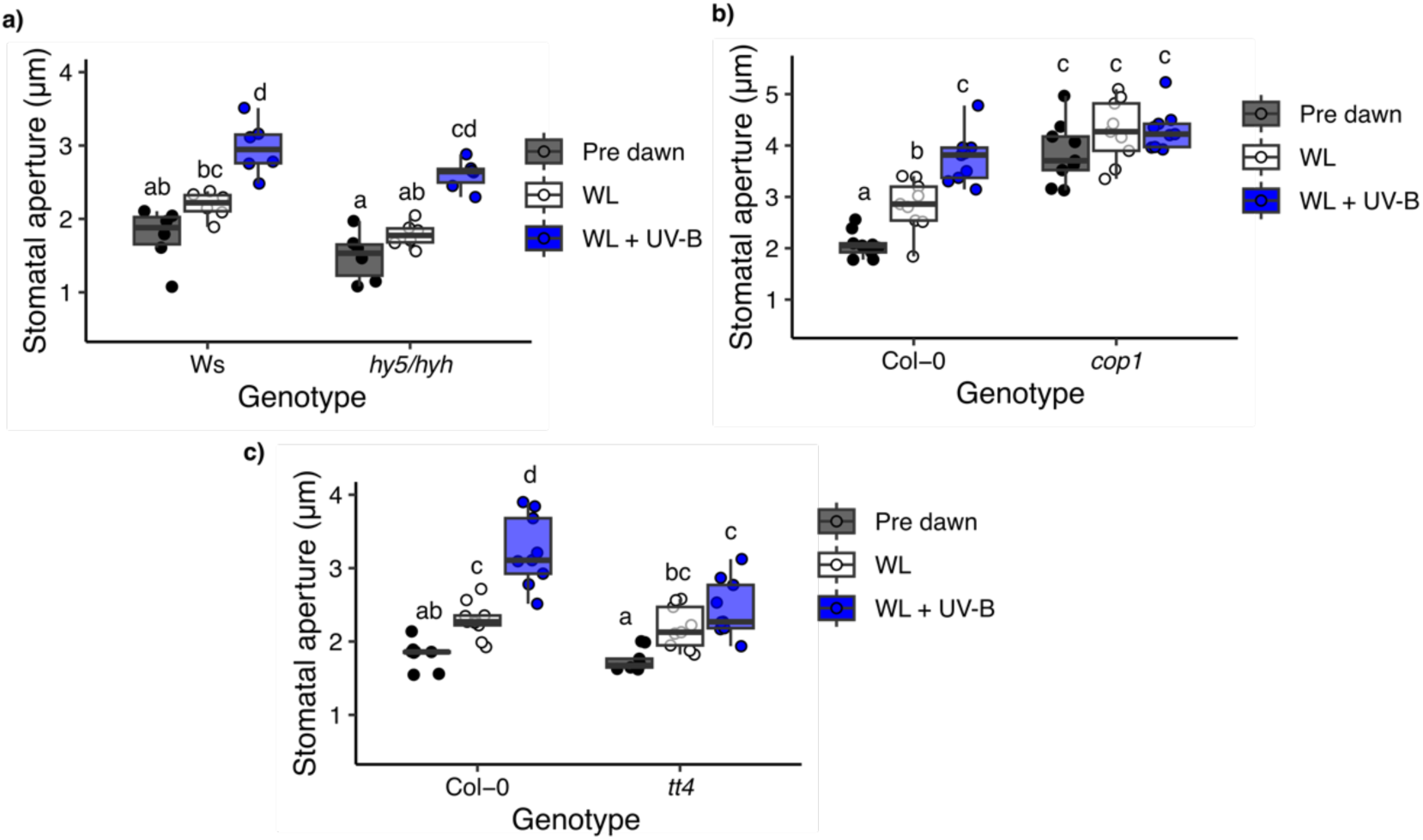
UV-B-mediated stomatal opening likely requires COP1, but not HY5/HYH. The stomatal apertures of **a)** *hy5hyh*, **b)** *cop1*, and **c)** *tt4* mutants following 6 h of WL ± UV-B treatment. Stomatal apertures were also measured prior to dawn (Pre dawn). Data are presented as boxplots showing the median and interquartile range of each group. The upper and lower whiskers represent data within 1.5 * IQR. Each individual plant’s mean stomatal aperture is represented as a point on the plot. For all genotype and treatment combinations, n = 9 seedlings over 3 independent experiments. Each mean seedling stomatal aperture was calculated from 10 stomatal measurements. Data were analysed using a 2-way ANOVA, followed by Tukey multiple comparison test. In **b)** *cop1* was grown in parallel with *uvr8-6* (Fig. 1C) so both plots use the same Col-0 control data.

**Fig. S4.**
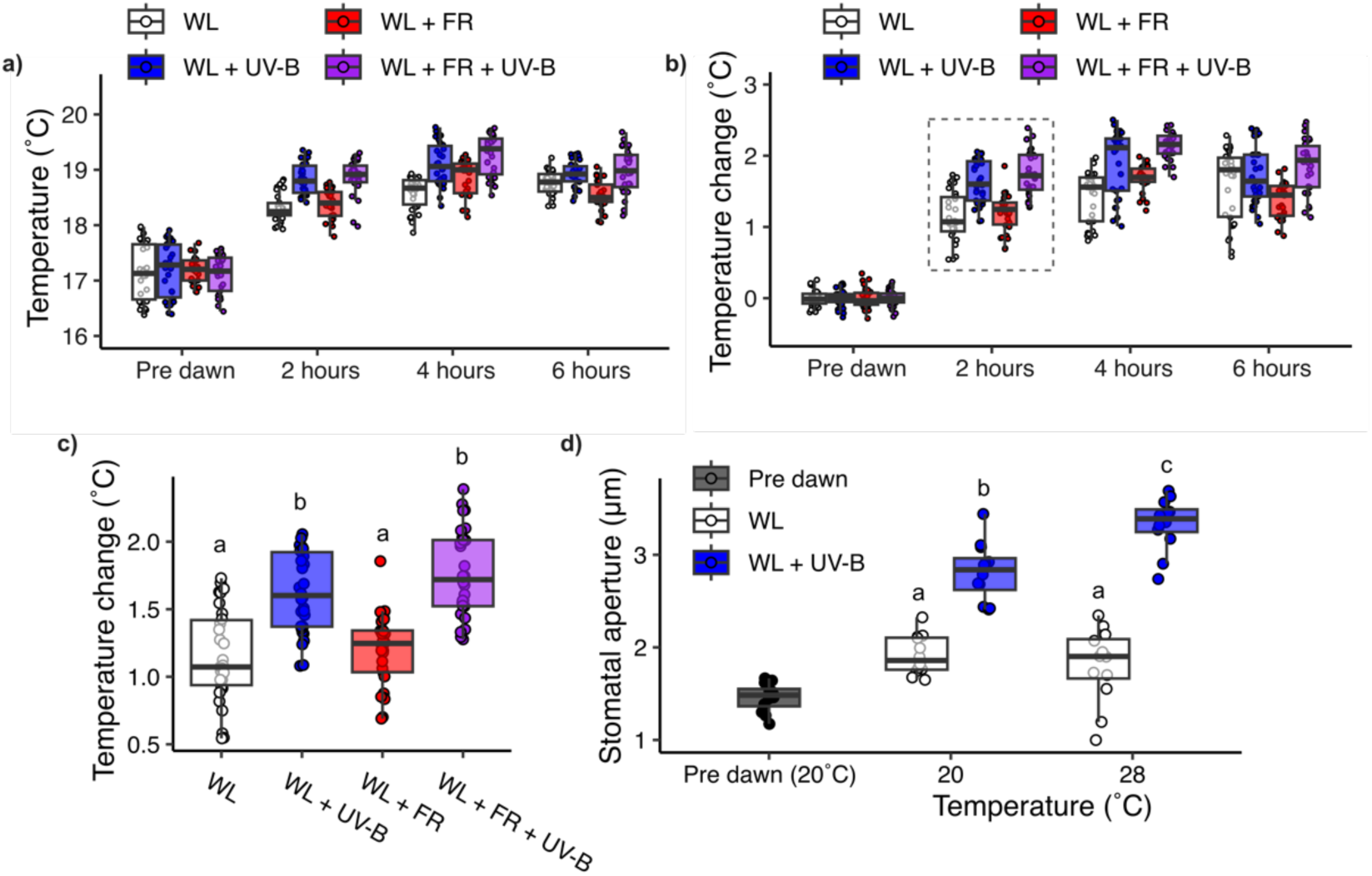
UV-B-mediated increases in stomatal aperture do not result from small elevations in cotyledon temperature. **a-c)** Cotyledon temperature measurements from 7 day old Col-0 seedlings treated with WL ± FR and/or UV-B from dawn. **a)** Absolute temperature measurements and **b)** Temperature change (relative to pre dawn values) over a 6 hour time-course. **c)** Temperature change at 2 hours (indicated by dashed box in **b**). **d)** Mean stomatal apertures of 7 day old Col-0 seedlings treated with WL ± UV-B at 20°C and 28°C for 6 hours following dawn. All data presented as points overlayed on top of boxplots showing the median and interquartile range (IQR) of each group. The upper and lower whiskers represent data within 1.5 * IQR. For **a-c)** all treatment combinations n = 10 seedlings over 3 independent experiments. For **d)** each mean stomatal aperture was calculated from 8-12 stomatal measurements. A total of n = 12 seedlings were analysed at each temperature treatment combination over 3 independent experiments. Data in **c)** was analysed using a 1-way ANOVA looking at the effect of light treatment, and in **d)** using a 2-way ANOVA looking at the effect of temperature and light treatment (pre-dawn values were not included in statistical analysis) followed by Tukey multiple comparison tests. Letters indicate significance at p < 0.05.

**Fig. S5.**
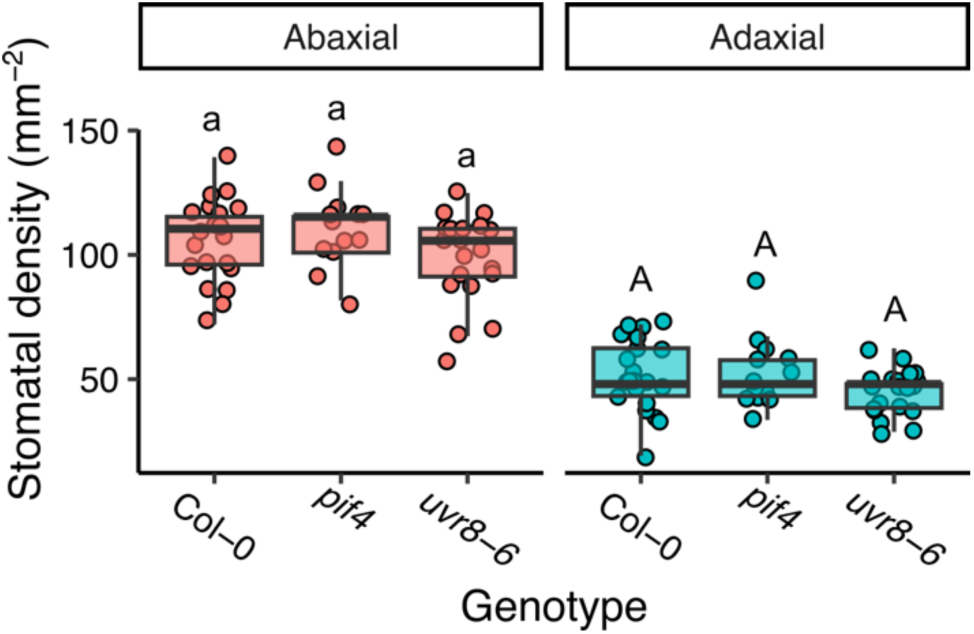
uvr8 and pif4 mutants show no significant differences in cotyledon stomatal density. The abaxial and adaxial stomatal densities of 7 day old Col-0, *pif4-101*, and *uvr8-6*. All data is presented as points overlayed on top of boxplots showing the median and interquartile range (IQR) of each group. The upper and lower whiskers represent data within 1.5 * IQR. n = 13-21 over 2 independent experiments. The abaxial and adaxial data in were separately analysed using a 1-way ANOVA. Multiple comparisons were performed using a post hoc Tukey test. Letters denote significance at p < 0.05.

**Fig. S6.**
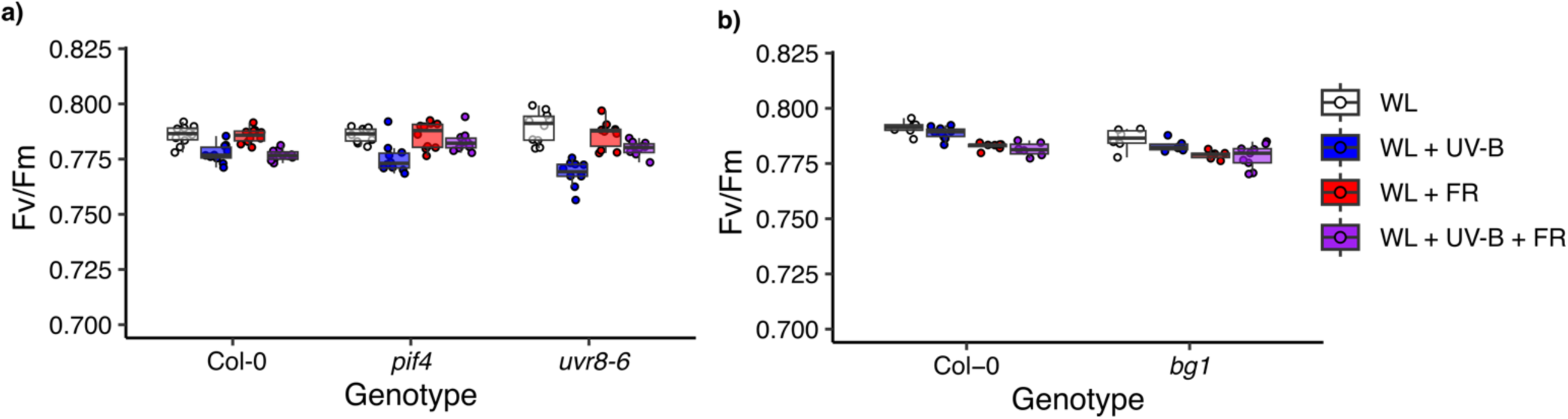
low dose UV-B supplementation has only minor effects on maximum photosystem II efficiency. **a-b)** F_V_/F_M_ measurements for 7 day old seedlings following 6 hours of WL ± FR ± UV-B treatment and 30 mins of dark adaption. Col-0 with **a)** *pif4-101* and *uvr8-6*, and **b)** *bg1* mutants are presented. Data in **a-b)** is one repeat representative of 3 independent experiments. Each experiment consists of n = 10 seedling F_V_/F_M_ measurements.

