## Supplementary figures for "Integration of light quality signals regulates ABA abundance and stomatal movements during seedling establishment"

### Supplementary Information

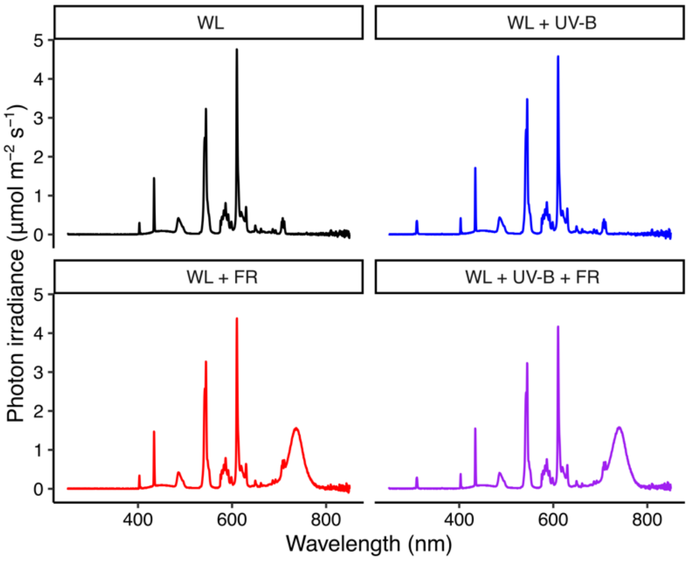


**Fig. S1 – Cabinet spectrums**

Light spectra of the different treatments described in this study. Every treatment includes ~75 µmolm$^{-2}$s$^{-1}$ white light (light between 400-700 nm). UV-B treatments involved the addition of 1 µmolm$^{-2}$s$^{-1}$ of UV-B light provided by a Philips TL100W/01 narrow band tube light. FR treatments involved the addition of FR LEDs with a maximum emission peak at 735 nm.

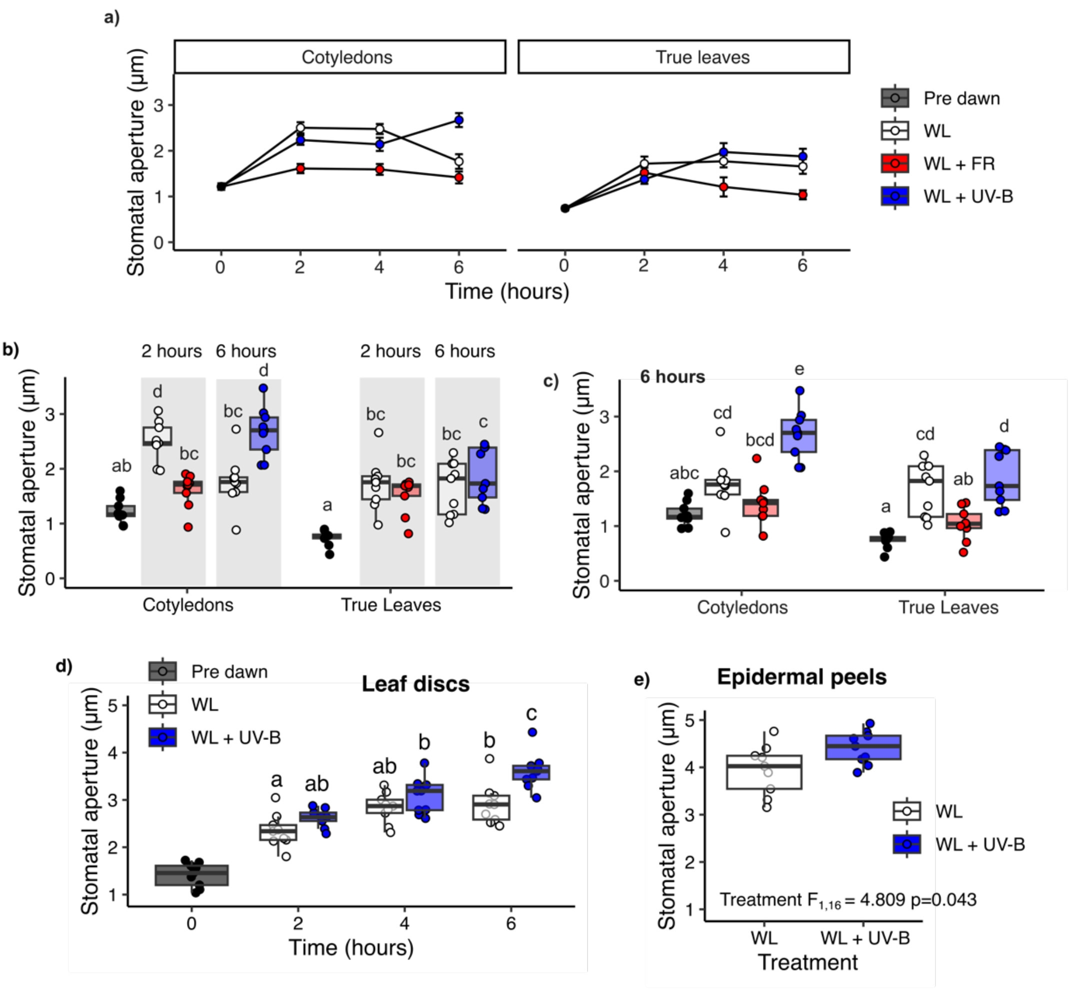


**Fig. S2 – Cotyledons and rosette leaves differ in stomatal responses to light quality**

**a)** Stomatal apertures from 7-day-old seedling cotyledons and 21-day-old true leaves were measured in WL, WL ± FR, and WL ± UV-B conditions over a 6 hour time course. Selected timepoints from **a)** are shown in **b)** 2h WL ± FR and 6h WL ± UV-B and **c)** all light treatments at 6 hours. Stomatal apertures from **d)** leaf discs and **e)** epidermal peels of 4-5 week-old Col-0 plants. Apertures were measured **d)** pre dawn and over a 6 h time course (2, 4, and 6 h) of WL ± UV-B treatment. In **e)** apertures were measured following 6 hours of WL ± UV-B treatment. For **a)** data is represented as mean ± s.e.m. For **b-e)**, data are presented as boxplots showing the median and interquartile range (IQR) of each group. The upper and lower whiskers represent data within 1.5 * IQR. Each individual plant's mean stomatal aperture calculated from 10-12 stomatal measurements is represented as a point on the plot. For all genotype treatment combinations, n = 9 plants over 3 independent experiments. Data in **b-d)** were analysed using 2-way ANOVA followed by Tukey Multiple Comparisons tests.

Data in **e)** was analysed using a Students T test. Letters indicate significance at $p < 0.05$.

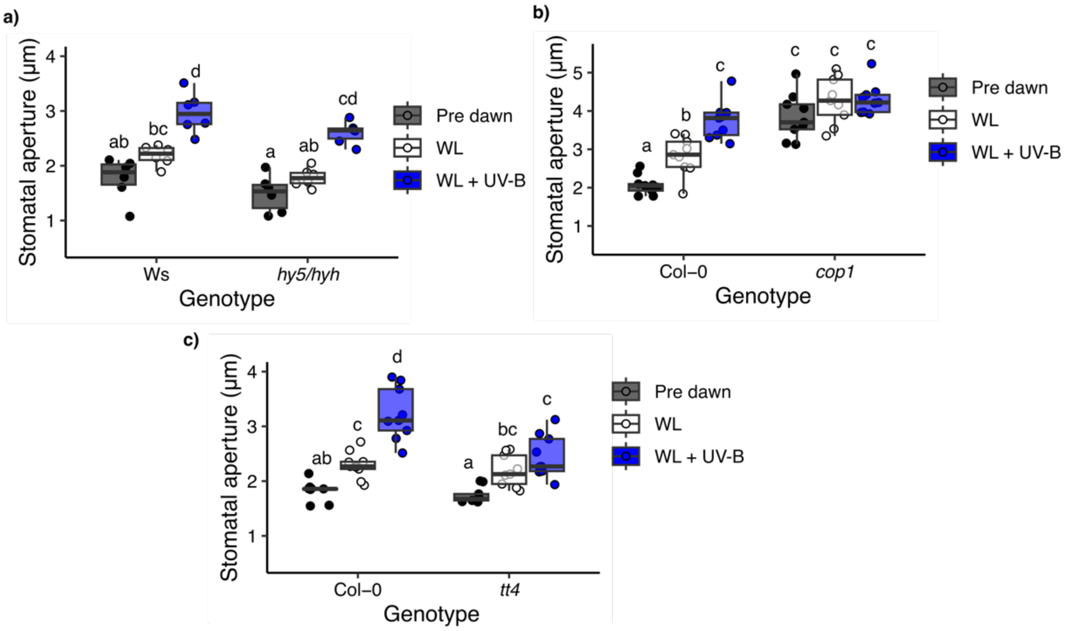


**Fig. S3 – UV-B-mediated stomatal opening likely requires COP1, but not HY5/HYH**

The stomatal apertures of **a)** *hy5hyh*, **b)** *cop1*, and **c)** *tt4* mutants following 6 h of WL ± UV-B treatment. Stomatal apertures were also measured prior to dawn (Pre dawn). Data are presented as boxplots showing the median and interquartile range of each group. The upper and lower whiskers represent data within 1.5 * IQR. Each individual plant's mean stomatal aperture is represented as a point on the plot. For all genotype and treatment combinations, n = 9 seedlings over 3 independent experiments. Each mean seedling stomatal aperture was calculated from 10 stomatal measurements. Data were analysed using a 2-way ANOVA, followed by Tukey multiple comparison test. In **b)** *cop1* was grown in parallel with *uvr8-6* (**Fig. 1C**) so both plots use the same Col-0 control data.

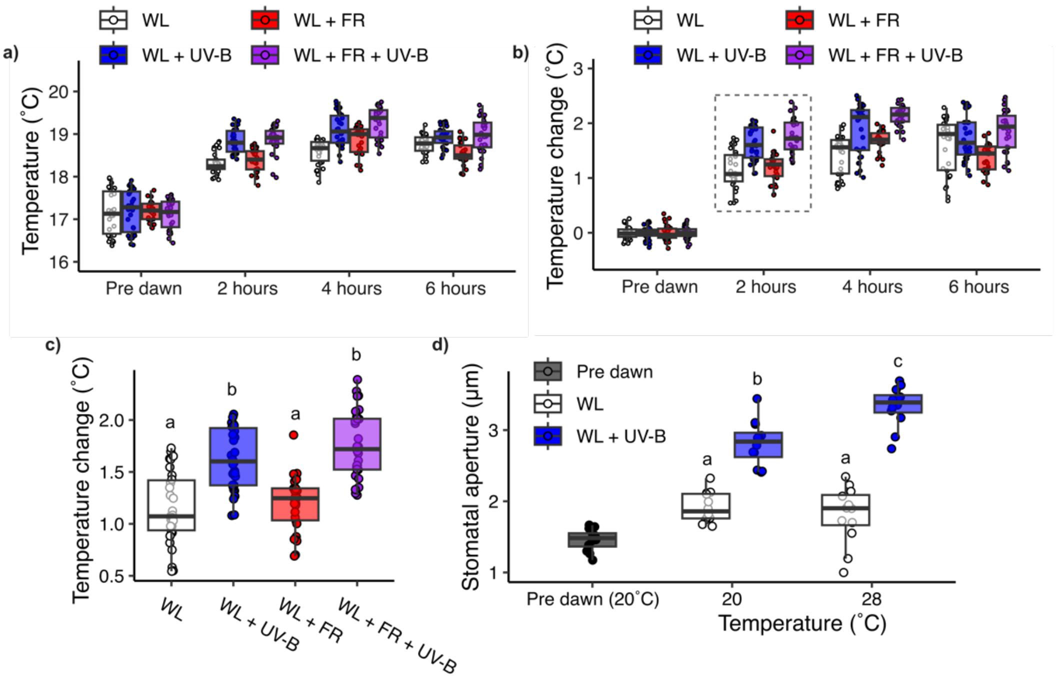


**Fig. S4 – UV-B-mediated increases in stomatal aperture do not result from small elevations in cotyledon temperature**

**a-c)** Cotyledon temperature measurements from 7 day old Col-0 seedlings treated with WL ± FR and/or UV-B from dawn. **a)** Absolute temperature measurements and **b)** Temperature change (relative to pre dawn values) over a 6 hour time-course. **c)** Temperature change at 2 hours (indicated by dashed box in **b**). **d)** Mean stomatal apertures of 7 day old Col-0 seedlings treated with WL ± UV-B at 20˚C and 28˚C for 6 hours following dawn. All data presented as points overlayed on top of boxplots showing the median and interquartile range (IQR) of each group. The upper and lower whiskers represent data within 1.5 * IQR. For **a-c)** all treatment combinations n = 10 seedlings over 3 independent experiments. For **d)** each mean stomatal aperture was calculated from 8-12 stomatal measurements. A total of n = 12 seedlings were analysed at each temperature treatment combination over 3 independent experiments. Data in **c)** was analysed using a 1-way ANOVA looking at the effect of light treatment, and in **d)** using a 2-way ANOVA looking at the effect of temperature and light treatment (pre-dawn values were not included in statistical analysis) followed by Tukey multiple comparison tests. Letters indicate significance at $p < 0.05$.

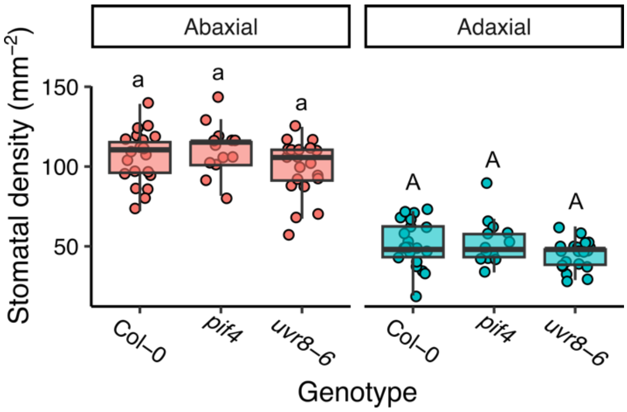


**Fig. S5 – uvr8 and pif4 mutants show no significant differences in cotyledon stomatal density**

The abaxial and adaxial stomatal densities of 7 day old Col-0, *pif4-101*, and *uvr8-6*. All data is presented as points overlayed on top of boxplots showing the median and interquartile range (IQR) of each group. The upper and lower whiskers represent data within 1.5 * IQR. n = 13-21 over 2 independent experiments. The abaxial and adaxial data in were separately analysed using a 1-way ANOVA. Multiple comparisons were performed using a post hoc Tukey test. Letters denote significance at $p < 0.05$.

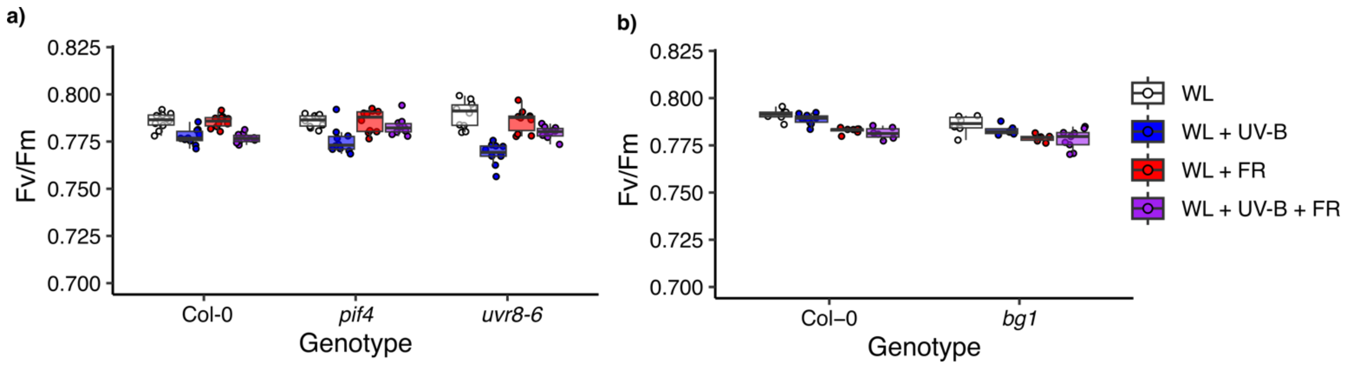


**Fig. S6 – low dose UV-B supplementation has only minor effects on maximum photosystem II efficiency**

**a-b)** $F_V/F_M$ measurements for 7 day old seedlings following 6 hours of WL ± FR ± UV-B treatment and 30 mins of dark adaption. Col-0 with **a)** *pif4-101* and *uvr8-6*, and **b)** *bg1* mutants are presented. Data in **a-b)** is one repeat representative of 3 independent experiments. Each experiment consists of n = 10 seedling $F_V/F_M$ measurements.
